# A chromosome-level genome assembly of the endangered northern dragonhead sap beetle, *Thymogethes norvegicus* (Easton, 1959)

**DOI:** 10.64898/2026.11.10.765633

**Authors:** Marius A. Strand, Ole K. Tørresen, Morten Skage, Giada Ferrari, Ave Tooming-Klunderud, Hallvard Elven, Kjetill S. Jakobsen

**Affiliations:** Department of Biosciences, University of Oslo, Oslo, Norway; Natural History Museum, University of Oslo, Oslo, Norway

**Keywords:** Earth Biogenome Project Norway, Nitidulidae, bacterial endosymbionts, chromosome-level genome assembly, conservation genomics, host-plant specialization

## Abstract

The northern dragonhead sap beetle (*Thymogethes norvegicus*) is believed to be endemic to Norway and is classified as endangered. Its life cycle is closely tied to the host plant northern dragonhead (*Dracocephalum ruyschiana*), itself classified as vulnerable in Norway. Here, we present a chromosome-level genome assembly of a male *T. norvegicus*. The assembly comprises two pseudo-haplotypes, hap1 and hap2, of 1,142 and 1,109 Mb, with 96.3% and 96.8% scaffolded into chromosomes, respectively. Hap1 contains 8 autosomes and the X chromosome, while hap2 contains 8 autosomes. Assembly quality was high, with conserved single-copy ortholog completeness scores of 96.6% and 90.7% and combined k-mer completeness of 98.1%. A total of 13,484 and 12,550 protein-coding genes were annotated in hap1 and hap2, respectively. This assembly provides a valuable genomic resource for the conservation of this endangered endemic species totally dependent on the vulnerable plant, the northern dragonhead.

## Introduction

The northern dragonhead sap beetle *Thymogethes norvegicus* (Easton, 1959) (Coleoptera: Nitidulidae) (Figure 1A,C) is believed to be endemic to Norway. Norway thus has a unique responsibility to protect it. The beetle is known from approximately fifty Norwegian localities, all within a radius of 60 km from the capital Oslo. This is also the most densely populated region in Norway, and the pressure on the beetle’s habitats is significant. The species is listed as Endangered (EN) in the Norwegian Red list (Artsdatabanken, 2021). The main threats are urban development, wear on biotopes in recreational areas, spread of invasive plant species, and changes in agricultural land use with intensification and abandonment. Recent surveys of the populations closest to the capital have shown that these are in steep decline (Elven, 2024b, 2024a). Out of 26 examined localities (about half of all known localities), almost half seem to have lost the beetle within just the last ten to fifteen years. Populations further from the capital are probably doing better, and several new populations have been discovered in recent years, including some that are quite large (Elven & Pavels, 2021; Olberg & Lønnve, 2021, 2022, 2024a, 2024b). However, both rural and urban populations face multiple pressures, and the decline is likely to continue.

**Figure 1.**
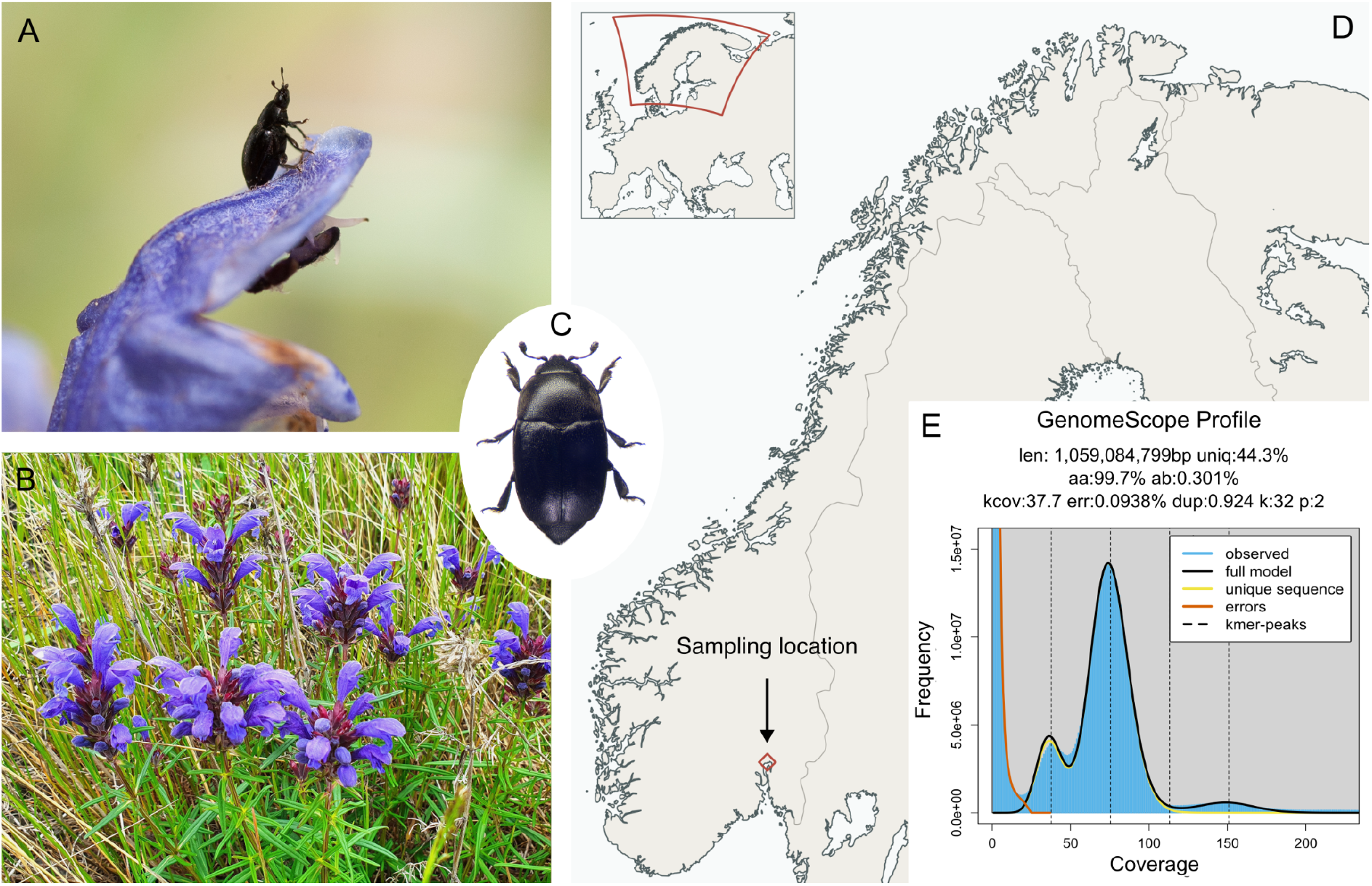
Thymogethes norvegicus, host plant, sampling locality, and genome profile. A) Adult *T. norvegicus* on *Dracocephalum ruyschiana*; photo by Hallvard Elven. **B)** Flowering *D. ruyschiana*, the host plant of *T. norvegicus*; photograph by Hallvard Elven. **C)** Dorsal view of *T. norvegicus*; photo by Hallvard Elven. **D)** Sampling locality in southern Norway (red diamond; 59.899591°N, 10.608603°E; coordinate uncertainty ±10 m). The main map shows Norway and parts of Sweden, Finland, and Russia. The inset shows Europe, with the red outline indicating the extent of the main map. Basemap data: Natural Earth 1:10m. **E)** GenomeScope profile of HiFi reads from the sequenced individual, estimating a genome size of 1,059 Mb and heterozygosity of 0.301%. The peaks at lower and higher k-mer coverage correspond to heterozygous and homozygous regions of the genome, respectively.

The larvae of the dragonhead beetle develop in the flowers of northern dragonhead *Dracocephalum ruyschiana L*. (Lamiaceae) (Figure 1B). The host plant is widely distributed but not common in continental parts of Southern Norway. It requires shallow, dry, alkaline soil and is typically found in dry meadows and pastures, as well as rock outcrops, from sea level up to the montane zone. The life cycle of the beetle has been studied by Hallvard Elven and Anders Endrestøl. The data are still unpublished, but a brief summary is given here: The eggs are laid in late May or early June, at the onset of the blooming of the dragonhead. The larvae develop within the flowers at the expense of the developing seeds. They finish their development in late June and leave the flowers to pupate in the ground. The pupae hatch in July, and the adults spend the autumn at the locality feeding on other available flowering plants. They hibernate in the ground, and mate and lay eggs in the following spring. The beetle has a far smaller distribution than the host plant and appears to have a much narrower climatic tolerance. Whereas the plant extends up to 800 meters above sea level, the beetle does not go above 250 meters.

The host plant is listed as Vulnerable in the Norwegian Red List (Artsdatabanken, 2021). Since 2010, the plant has had status as a Prioritized Species with its own action plan and annual funding for conservation (Direktoratet for naturforvaltning, 2010). It is juridically protected against all damage, and the protection also extends to its habitat. The dragonhead beetle is covered by the same action plan and likewise receives annual funding for conservation, but it does not have status as a Prioritized Species and has no juridical protection. Several conservation measures have been carried out since the plant became a Prioritized Species. These include surveying, habitat protection, habitat management/restoration, and – in the case of the beetle – re-introduction to former localities and a pilot project on captive breeding. Despite these efforts, a status report from 2024 concluded that, while our knowledge about the two species has increased significantly since 2010, 13 years of conservation measures have not led to a positive change in red list status for either species (Røsok, 2024).

As existing localities continue to degrade and go extinct, future conservation measures for the dragonhead beetle will probably increasingly involve *ex situ* breeding and relocation. In this context, having more knowledge about the genetics of the species will be highly useful for making informed decisions about e.g. which localities to harvest individuals from and where to release them. A high-quality genome assembly will greatly facilitate future genetic studies.

Motivated by the delicate dependency of the dragonhead sap beetle and its host plant, and the vulnerability of this interaction from a conservation biology angle, we have generated a chromosome level reference genome of a male beetle using long-read sequencing and chromosomal scaffolding. The genome will be an important resource in conservation efforts for this endangered beetle. As a completion of the needed reference genome resources, in a companion paper (Strand et al., 2026), we present a chromosome level assembly of the host plant, northern dragonhead as well.

The dragonhead sap beetle genome assembly was generated as part of the Earth BioGenome Project Norway.

## Material and Methods

### Sample acquisition and DNA extraction

Two male adult specimens of *T. norvegicus* were collected on the host plant *D. ruyschiana* at Oksenøya Bruk (Norway, Akershus, Bærum; 59.899591°N, 10.608603°E ± 10 m; Figure 1D) on May 22, 2024. The locality houses one of the largest remaining populations of the dragonhead beetle close to the capital Oslo. No permit was needed for collecting the material. The specimens were frozen alive using liquid nitrogen. Prior to freezing, the genitalia and a front leg from each specimen were secured as a voucher. The material was stored at minus 80 ^o^C. The vouchers are deposited at the Natural History Museum, University of Oslo.

DNA isolation for PacBio long-read sequencing started from the remaining body of one specimen following the Sanger Tree of Life manual MagAttract v3 HMW DNA extraction protocol (Oatley et al., 2025). The DNA was eluted in a volume of 300 μL and the final SPRI step was omitted. Quality check of the amount, purity and integrity of the isolated DNA was performed using a combination of Qubit HS DNA quantification assay kit (Thermo Fisher) and Fragment Analyzer (DNA HS 50 kb large fragment kit, Agilent Tech.).

### Library preparation and sequencing for *de novo* assembly

All HMW DNA isolated from a single beetle was sheared into an average fragment size of 15-20 kb large fragments using the Hamilton Microlab Prep instrument (Hamilton). A HiFi library was prepared following the PacBio protocol for HiFi library preparation using the SMRTbell® Prep Kit 3.0. The final HiFi library was size-selected with a 3 kb cut-off using Ampure-PB beads and sequencing was performed by the Norwegian Sequencing Centre on a PacBio Revio instrument (Pacific Biosciences Inc). The library was sequenced on a 25M SMRT cell using SPRQ Polymerase kit and sequencing plate.

A Hi-C library was prepared using the Arima High Coverage HiC kit (Arima Genomics), following the manufacturer’s recommendations for low input samples and starting with the entire specimen. Final library quality was quantified using a Kapa Library quantification kit for Illumina (Roche Inc.). The library was sequenced with other libraries on an Illumina NovaSeqX 25B flowcell (Illumina Inc) with 2*150 bp paired end mode at the Norwegian Sequencing Centre (https://www.sequencing.uio.no).

### Genome assembly and curation, annotation and evaluation

A full list of relevant software tools and versions is presented in Table 1. We assembled the species using a pre-release of the Earth BioGenome Project Norway (EBP-Nor) genome assembly pipeline (https://github.com/ebp-nor/GenomeAssembly). The main steps of this pipeline are described below. KMC (Kokot et al., 2017) was used to count k-mers of size 32 in the PacBio HiFi reads, excluding k-mers occurring more than 10,000 times. GenomeScope (Ranallo-Benavidez et al., 2020) was run as part of the pipeline on the k-mer histogram output from KMC to estimate genome size, heterozygosity and repeat content (Figure 1E). Ploidy level was investigated using Smudgeplot (Ranallo-Benavidez et al., 2020). HiFiAdapterFilt (Sim et al., 2022) was applied on the HiFi reads to remove possible remnant PacBio adapter sequences. The filtered HiFi reads were assembled using hifiasm (Cheng et al., 2021) with Hi-C integration resulting in a pair of haplotype-resolved assemblies, pseudo-haplotype one (hap1) and pseudo-haplotype two (hap2). Unique k-mers in each assembly/pseudo-haplotype were identified using meryl (Rhie et al., 2020) and used to create two sets of Hi-C reads, one without any k-mers occurring uniquely in hap1 and the other without k-mers occurring uniquely in hap2.

**Table 1.**
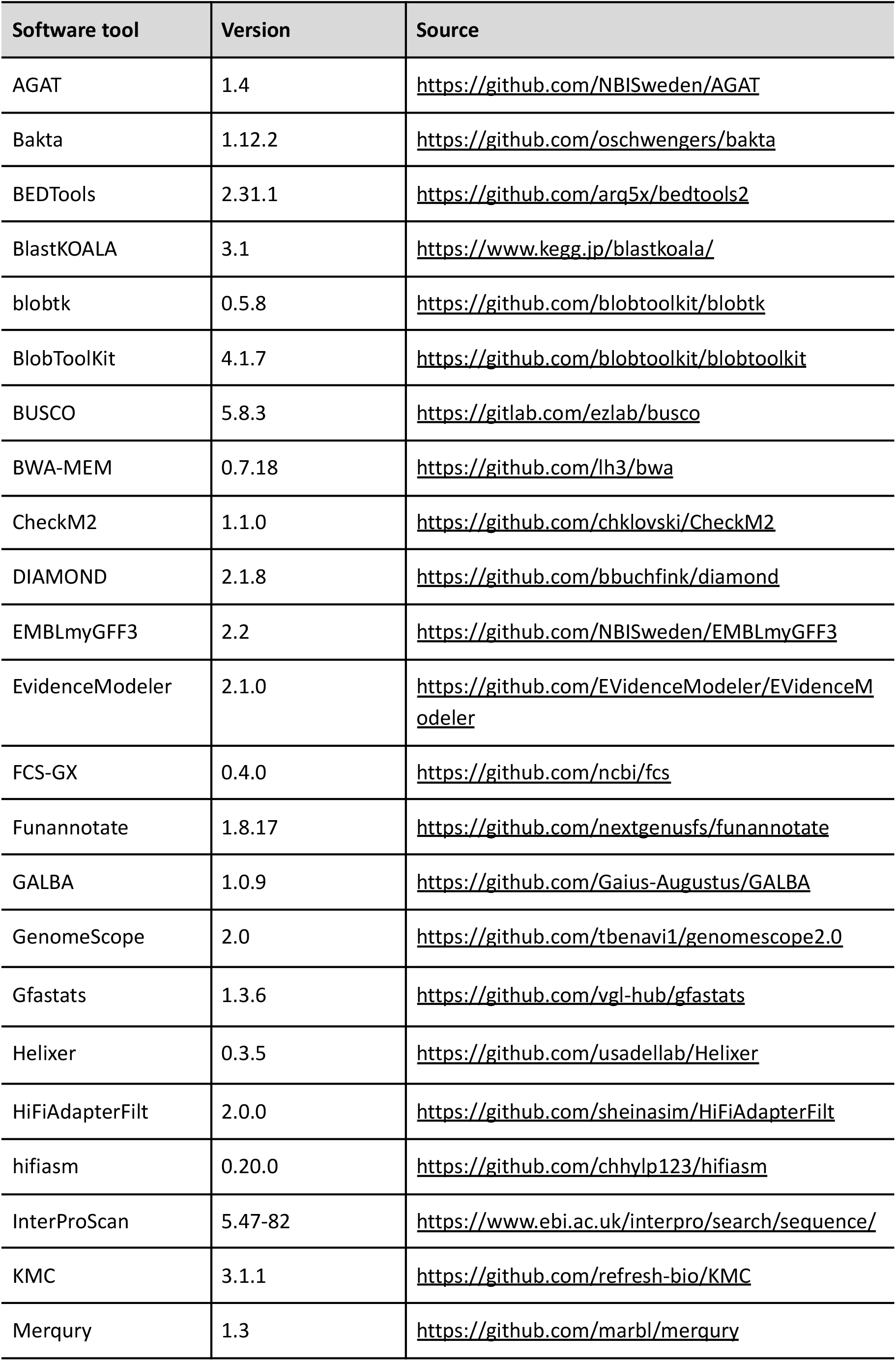
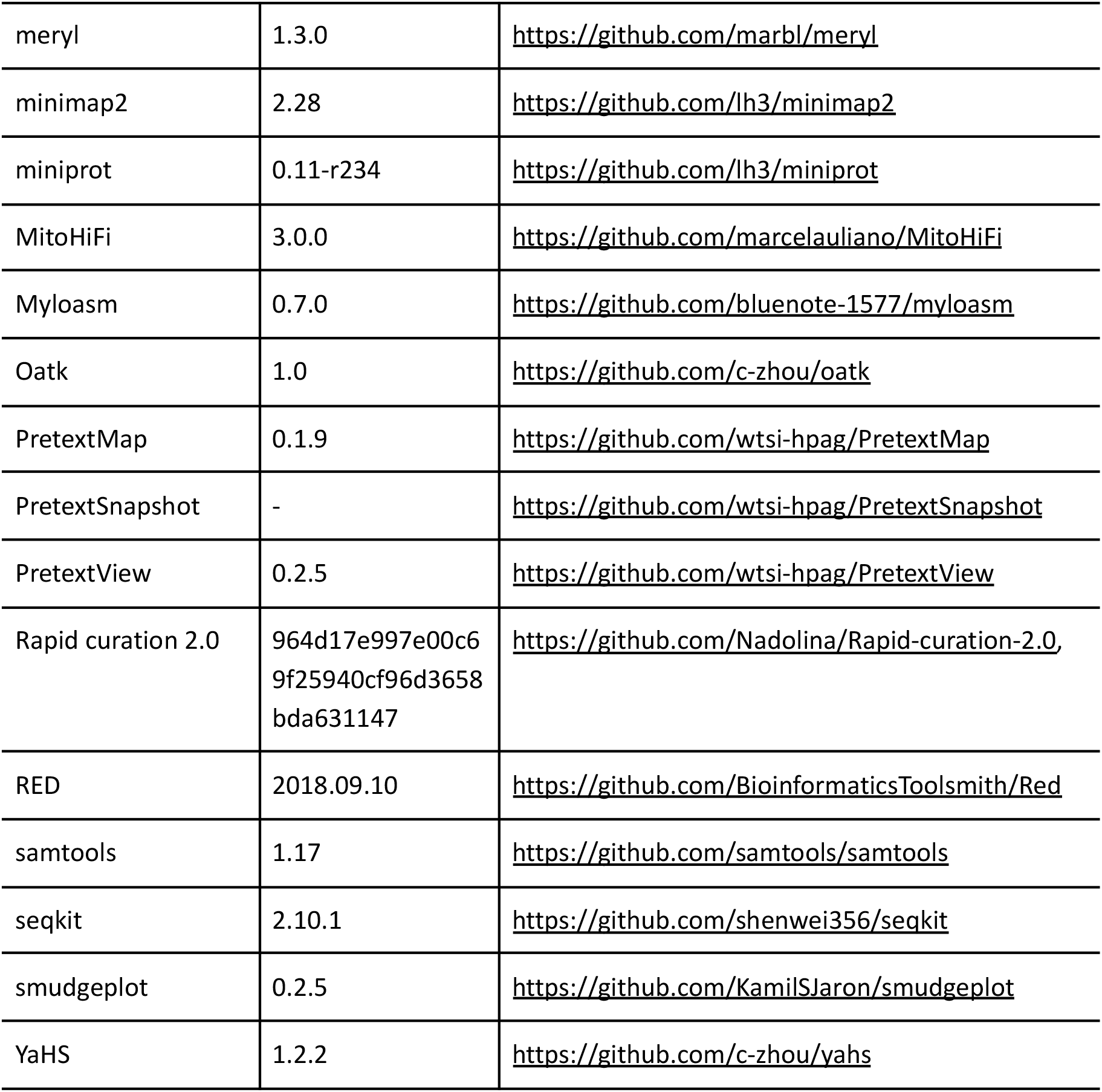
Software tools: versions and sources.

| **Software tool** | **Version** | **Source** |
|---|---|---|
| AGAT | 1.4 | https://github.com/NBISweden/AGAT |
| Bakta | 1.12.2 | https://github.com/oschwengers/bakta |
| BEDTools | 2.31.1 | https://github.com/arq5x/bedtools2 |
| BlastKOALA | 3.1 | https://www.kegg.jp/blastkoala/ |
| blobtk | 0.5.8 | https://github.com/blobtoolkit/blobtk |
| BlobToolKit | 4.1.7 | https://github.com/blobtoolkit/blobtoolkit |
| BUSCO | 5.8.3 | https://gitlab.com/ezlab/busco |
| BWA-MEM | 0.7.18 | https://github.com/lh3/bwa |
| CheckM2 | 1.1.0 | https://github.com/chklovski/CheckM2 |
| DIAMOND | 2.1.8 | https://github.com/bbuchfink/diamond |
| EMBLmyGFF3 | 2.2 | https://github.com/NBISweden/EMBLmyGFF3 |
| EvidenceModeler | 2.1.0 | https://github.com/EVidenceModeler/EVidenceModeler |
| FCS-GX | 0.4.0 | https://github.com/ncbi/fcs |
| Funannotate | 1.8.17 | https://github.com/nextgenusfs/funannotate |
| GALBA | 1.0.9 | https://github.com/Gaius-Augustus/GALBA |
| GenomeScope | 2.0 | https://github.com/tbenavi1/genomescope2.0 |
| Gfastats | 1.3.6 | https://github.com/vgl-hub/gfastats |
| Helixer | 0.3.5 | https://github.com/usadellab/Helixer |
| HiFiAdapterFilt | 2.0.0 | https://github.com/sheinasim/HiFiAdapterFilt |
| hifiasm | 0.20.0 | https://github.com/chhylp123/hifiasm |
| InterProScan | 5.47-82 | https://www.ebi.ac.uk/interpro/search/sequence/ |
| KMC | 3.1.1 | https://github.com/refresh-bio/KMC |
| Merqury | 1.3 | https://github.com/marbl/merqury |
| meryl | 1.3.0 | https://github.com/marbl/meryl |
| minimap2 | 2.28 | https://github.com/lh3/minimap2 |
| miniprot | 0.11-r234 | https://github.com/lh3/miniprot |
| MitoHiFi | 3.0.0 | https://github.com/marcelauliano/MitoHiFi |
| Myloasm | 0.7.0 | https://github.com/bluenote-1577/myloasm |
| Oatk | 1.0 | https://github.com/c-zhou/oatk |
| PretextMap | 0.1.9 | https://github.com/wtsi-hpag/PretextMap |
| PretextSnapshot | - | https://github.com/wtsi-hpag/PretextSnapshot |
| PretextView | 0.2.5 | https://github.com/wtsi-hpag/PretextView |
| Rapid curation 2.0 | 964d17e997e00c69f25940cf96d3658bda631147 | https://github.com/Nadolina/Rapid-curation-2.0, |
| RED | 2018.09.10 | https://github.com/BioinformaticsToolsmith/Red |
| samtools | 1.17 | https://github.com/samtools/samtools |
| seqkit | 2.10.1 | https://github.com/shenwei356/seqkit |
| smudgeplot | 0.2.5 | https://github.com/KamilSJaron/smudgeplot |
| YaHS | 1.2.2 | https://github.com/c-zhou/yahs |

K-mer filtered Hi-C reads were aligned to each scaffolded assembly using BWA-MEM (Li, 2013) with -5SPM options. The alignments were sorted based on name using samtools (Li et al., 2009) before applying samtools fixmate to remove unmapped reads and secondary alignments and to add mate score, and samtools markdup to remove duplicates. The resulting BAM files were used to scaffold the two assemblies using YaHS (Zhou et al., 2023) with default options. FCS-GX (Astashyn et al., 2024) was used to search for putative contamination and contaminated sequences were removed. The mitochondrion was searched for in contigs and reads using Oatk (Zhou et al., 2025). The assemblies were manually curated using PretextView and Rapid curation 2.0. Chromosomes were identified inspecting the Hi-C contact map in PretextView.

We annotated the genome assemblies using a pre-release version of the EBP-Nor genome annotation pipeline (https://github.com/ebp-nor/GenomeAnnotation). The main steps of this pipeline are described below. First, AGAT (https://zenodo.org/record/7255559) agat_sp_keep_longest_isoform.pl and agat_sp_extract_sequences.pl were used on the *Drosophila melanogaster* (BDGP6.46, Ensembl release 110) genome assembly and annotation to generate one protein (the longest isoform) per gene. Miniprot (Li, 2023) was used to align the proteins to the curated assemblies. UniProtKB/Swiss-Prot (UniProt Consortium, 2023) release 2025_03 in addition to the Arthropoda part of OrthoDB v12 (Tegenfeldt et al., 2025) were also aligned separately to the assemblies. RED (Girgis, 2015) was run via redmask (https://github.com/nextgenusfs/redmask) on the assemblies to mask repetitive areas. In addition to gene models from protein alignments, GALBA (Brůna et al., 2023; Buchfink et al., 2015; Hoff & Stanke, 2019; Li, 2023; Stanke et al., 2006) was run with the *Drosophila melanogaster* proteins using the miniprot mode on the masked assemblies to generate *ab initio* predicted gene models. Helixer (Holst et al., 2025) was run using the invertebrate-specific model (invertebrate_v0.3_m_0100.h5).

The funannotate-runEVM.py script from Funannotate was used to run EvidenceModeler (Haas et al., 2008) on the alignments of BDGP6.46 proteins, UniProtKB/Swiss-Prot proteins, Arthropoda proteins and the predicted genes from GALBA and Helixer. The resulting predicted proteins were compared to the protein repeats that Funannotate distributes using DIAMOND blastp and the predicted genes were filtered based on this comparison using AGAT. The filtered proteins were compared to the UniProtKB/Swiss-Prot release 2025_03 using DIAMOND (Buchfink et al., 2015) blastp to find gene names and InterProScan (Jones et al., 2014) was used to discover functional domains. AGATs agat_sp_manage_functional_annotation.pl was used to attach the gene names and functional annotations to the predicted genes. EMBLmyGFF3 (Norling et al., 2018) was used to combine the FASTA files and GFF3 files into a EMBL format for submission to ENA.

All the evaluation tools have also been implemented in a pipeline, similar to assembly and annotation (https://github.com/ebp-nor/GenomeEvaluation). The main steps of this pipeline are described below. Merqury (Rhie et al., 2020) was used to assess the completeness and quality of the genome assemblies by comparing them to the k-mer content of both the Hi-C reads and PacBio HiFi reads. BUSCO (Manni et al., 2021) was used to assess the completeness of the genome assemblies by comparing against the expected gene content in the polyphaga lineage (odb12.2). Gfastats (Formenti et al., 2022) was used to output different assembly statistics of the assemblies.

BlobToolKit and BlobTools2 (Laetsch & Blaxter, 2017), in addition to blobtk were used to visualize assembly statistics. To generate the Hi-C contact map image, the Hi-C reads were mapped to the assemblies using BWA-MEM (Li, 2013) using the same approach as above. Finally, PretextMap was used to create a contact map which was visualized using PretextSnapshot.

To characterize the differences between the two pseudo-haplotypes, we ran a genome alignment using minimap2 (Li, 2018) on the homologous chromosomes. The resulting alignment was processed with the minimap2-included paftools.js producing a report listing the number of insertions, SNPs and indels between the two pseudo-haplotypes.

The text in Methods and parts of Results is based on a template we use for all the species we publish in the EBP-Nor project.

### Evaluation of putative endosymbionts

Bacterial scaffolds identified by FCS-GX (Astashyn et al., 2024) in both haplotypes were evaluated alongside assemblies generated independently using Myloasm (Shaw et al., 2026). For the latter, HiFi reads were screened against 31-mers from the chromosome-scale beetle scaffolds, retaining reads with fewer than 50% matching k-mer positions. Candidate bacterial genomes were compared with each other using minimap2 (Li, 2018) and evaluated with CheckM2 (Chklovski et al., 2023). Circularity was assessed by mapping the original HiFi reads using minimap2 to the candidate genomes and to versions rotated by half their length using SeqKit (Shen et al., 2024). One genome was circularized by removing a duplicated terminal overlap. Taxonomic identification was performed using NCBI BLASTN searches of 16S rRNA genes (Zhang et al., 2000), with *ftsZ* providing additional support for *Wolbachia* supergroup assignment (Wang et al., 2020; Werren et al., 1995). The final circular genomes comprised *Wolbachia* from an FCS-GX-identified scaffold and *Symbiodolus* from the Myloasm assembly and were annotated using Bakta (Schwengers et al., 2021). Metabolic pathways were investigated using BlastKOALA (Kanehisa et al., 2016) and visualized via KEGG Mapper (Kanehisa et al., 2022).

## Results

### *De novo* genome assembly and annotation

The genome of *Thymogethes norvegicus* was assembled from a total of 74-fold coverage in Pacific Biosciences single-molecule HiFi long reads and 54-fold coverage in Arima Hi-C reads resulting in two haplotype-separated assemblies. The final assemblies have total lengths of 1142 Mb and 1109 Mb (Table 2 and Figure 2), respectively. Pseudo-haplotypes (hereafter termed haplotypes) one (hap1) and two (hap2) have scaffold N50 size of 172.3 Mb and 169.7 Mb, respectively, and contig N50 of 3.0 Mb and 2.8 Mb, respectively (Table 2 and Fig. 2). 8 pseudo-chromosomes were identified in both haplotypes, in addition to the X sex chromosome in hap1. The autosomes were named according to decreasing size.

**Table 2.** Genome data for *Thymogethes norvegicus*.

| **Project accession data** |  |  |
|---|---|---|
| Species | *Thymogethes norvegicus* |  |
| Specimen | HiFi from icThyNorv2 (Hi-C from icThyNorv1) |  |
| NCBI Taxonomy ID | 1974573 |  |
| BioProject | PRJEB129340 |  |
| BioSample ID | icThyNorv2: SAMEA124439599, icThyNorv1: SAMEA124437175 |  |
| Isolate information | Entire specimen |  |
| **Raw data accessions** |  |  |
| PacBio HiFi reads | ERR18092818 | PACBIO_SMRT (Revio) run: 9.1 M reads, 84.16 Gb |
| Hi-C Illumina reads | ERR18092792 and ERR18092796 | 2 ILLUMINA (Illumina NovaSeqX 25B) run: 204 M pairs of reads, 61.4 Gb |
| **Genome assembly metrics** |  |  |
| HiFi read coverage | 74x |  |
| Assembly | hap1 | hap2 |
| Assembly accession | GCA_990133344 | GCA_990133343 |
| Assembly identifier | icThyNorv2.1.hap1 | icThyNorv2.1.hap2 |
| Span (Mb) | 1142 | 1109 |
| Number of Contigs | 1378 | 1082 |
| Contig N50 length (Mb) | 3.0 | 2.8 |
| Longest contig (Mb) | 9.8 | 9.7 |
| Number of gaps | 715 | 710 |
| Number of Scaffolds | 667 | 375 |
| Scaffold N50 length (Mb) | 172.3 | 169.7 |
| Longest scaffold (Mb) | 222.4 | 224.7 |
| Percentage of assembly mapped to chromosomes | 96.3 | 96.8 |
| Sex chromosomes | X |  |
| Organelles | MT |  |
| Consensus quality (QV) compared to |  |  |
| Hi-C | 32.17 | 31.34 |
| HiFi | 54.82 | 54.93 |
| K-mer completeness (%) compared to |  |  |
| Hi-C | 96.3 | 96.1 |
| Hi-C both | 96.4 |  |
| HiFi | 94.1 | 91.4 |
| HiFi both | 98.1 |  |
| **Assembly comparison** (hap2 aligned to hap1) |  |  |
| Bases in alignment | 979,664,704 |  |
| Substitutions (%) | 649,692 (0.07) |  |
| 1bp deletions | 52,863 |  |
| 1bp insertions | 52,419 |  |
| 2bp deletions | 19,724 |  |
| 2bp insertions | 19,690 |  |
| [3,50) deletions | 75,420 |  |
| [3,50) insertions | 75,874 |  |
| [50,1000) deletions | 12,555 |  |
| [50,1000) insertions | 12,405 |  |
| >=1000 deletions | 3,234 |  |
| >=1000 insertions | 3,100 |  |
| **Genome annotation** |  |  |
| Assembly | hap1 | hap2 |
| Number of protein-coding genes | 13,484 | 12,550 |
| Number of protein-coding genes with functional domain | 12,484 | 11,615 |
| Number of protein-coding genes with gene names | 7,763 | 7,186 |
| BUSCO** | C:96.6%[S:94.8%,D:1.8%],F:2.2%,M:1.2%,n:3542 | C:90.7%[S:88.8%,D:1.8%],F:2.3%,M:7.1%,n:3542 |

\*Number of genes annotated with a functional domain as found by InterProScan. \*\* BUSCO scores based on the polyphaga\_odb12.2 BUSCO set using v5.8.3. C = complete [S = single copy, D = duplicated], F = fragmented, M = missing, n = number of orthologues in comparison.

**Figure 2.**
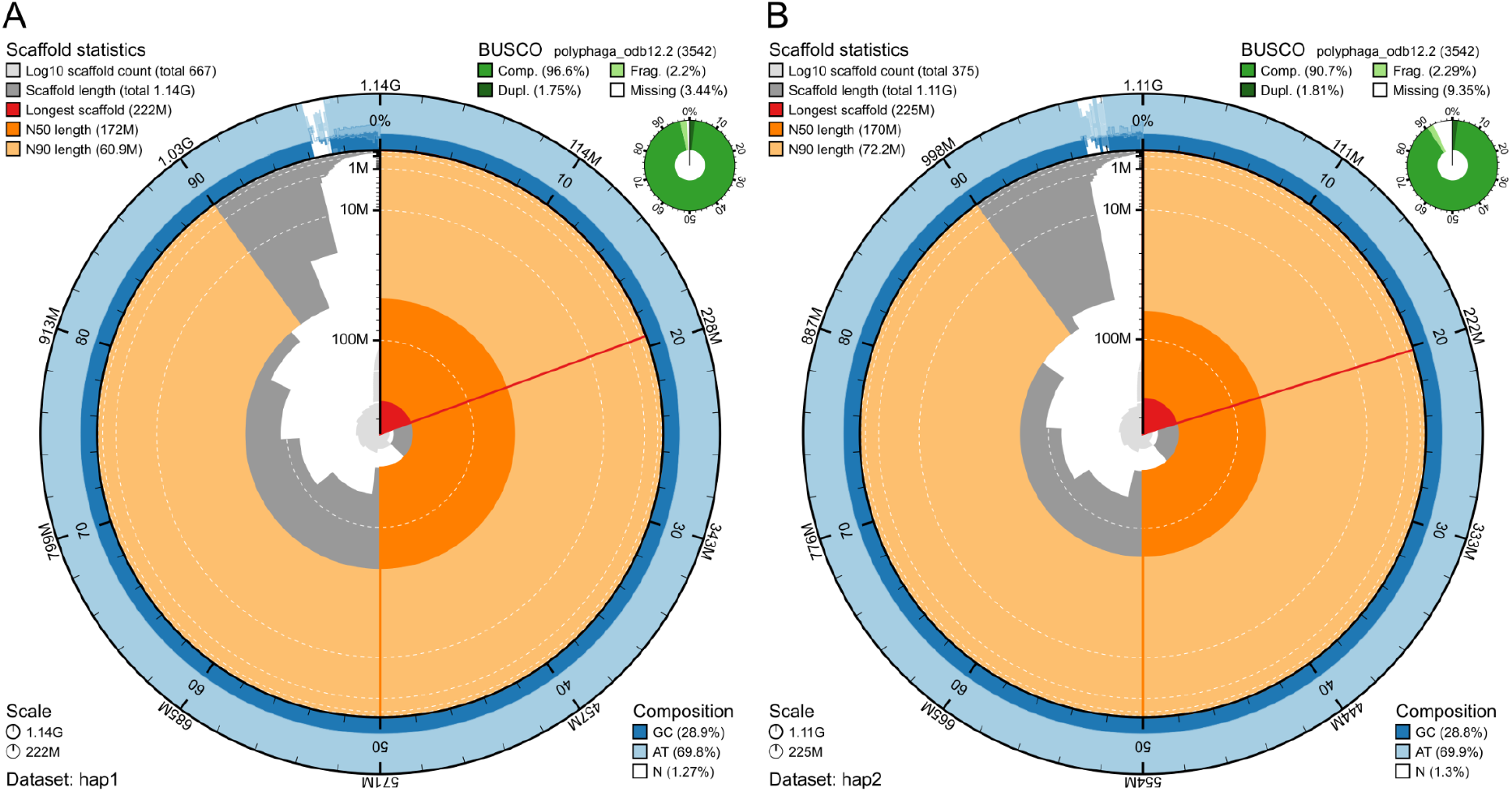
Metrics of the genome assemblies of Thymogethes norvegicus hap1 and hap2. A) Hap1. B) Hap2. The BlobToolKit Snailplots show N50 metrics and BUSCO gene completeness. The two outermost bands of the circle signify GC versus AT composition at 0.1% intervals, with mean, maximum, and minimum. The third outermost shows the N90 scaffold length, while the fourth is N50 scaffold length. The line from middle to second outermost band shows the size of the largest scaffold. All the scaffolds are arranged in a clockwise manner from largest to smallest, and shown in darker gray with white lines at different orders of magnitude, shown as a scale on the radius. The light gray shows the cumulative scaffold count. The scale inset in the lower left corner shows the total amount of sequence in the whole circle, and the fraction of the circle encompassed in the largest scaffold.

The Hi-C contact maps show well-resolved chromosome-scale scaffolds in both pseudo-haplotypes (Supplementary Figure 1).

When compared to a k-mer database of the Hi-C reads, hap1 had a k-mer completeness of 96.3%, hap2 of 96.1%, and combined they have a completeness of 96.4% (Table 2). Further, hap1 had an assembly consensus quality value (QV) of 32.17 and hap2 of 31.34, where a QV of 40 corresponds to one error every 10,000 bp, or 99.99% accuracy compared to a k-mer database of the Hi-C reads (QV 54.8 and 54.9, respectively, compared to a k-mer database of the HiFi reads) (Table 2). Merqury k-mer spectra showed that most assembly k-mers were assigned to the expected shared or haplotype-specific categories, supporting the quality of the haplotype-resolved assembly (Supplementary Fig. 2). When comparing the two pseudo-haplotypes using minimap2, there are 649,692 SNP differences (0.07% of the aligned sequence), 163,796 deletions in hap2 compared to hap1 ranging from 1 bp to more than 1000 bp and 163,488 insertions from 1 bp to more than 1000 bp in size. A total of 13484 and 12550 protein-coding genes were annotated in hap1 and hap2, respectively (Table 2).

Two bacterial endosymbionts were recovered, with 16S rRNA sequences closely matching *Wolbachia* (99.8% identity; OZ035014.1) and *Candidatus Symbiodolus clandestinus* (99.81%; PQ200766.1). The *ftsZ* (filamenting temperature-sensitive mutant Z) gene supported assignment of *Wolbachia* to supergroup A (100%; OZ035014.1). The selected circular genomes were 1.74 Mb (*Wolbachia*) and 1.49 Mb (*Symbiodolus*) (Figure 3), with CheckM2 completeness estimates of 98.83% and 100%, respectively. Both had 100% HiFi read coverage breadth, with 352 and 123 reads spanning their circular junctions. The *Symbiodolus* genome contained two identical 16S rRNA genes, each within a 16S–23S–5S rRNA gene cluster. An additional 86.7 kb circular, transposase-rich element, supported by 263 junction-spanning reads and approximately threefold higher read coverage, may represent an associated *Symbiodolus* plasmid.

**Figure 3.**
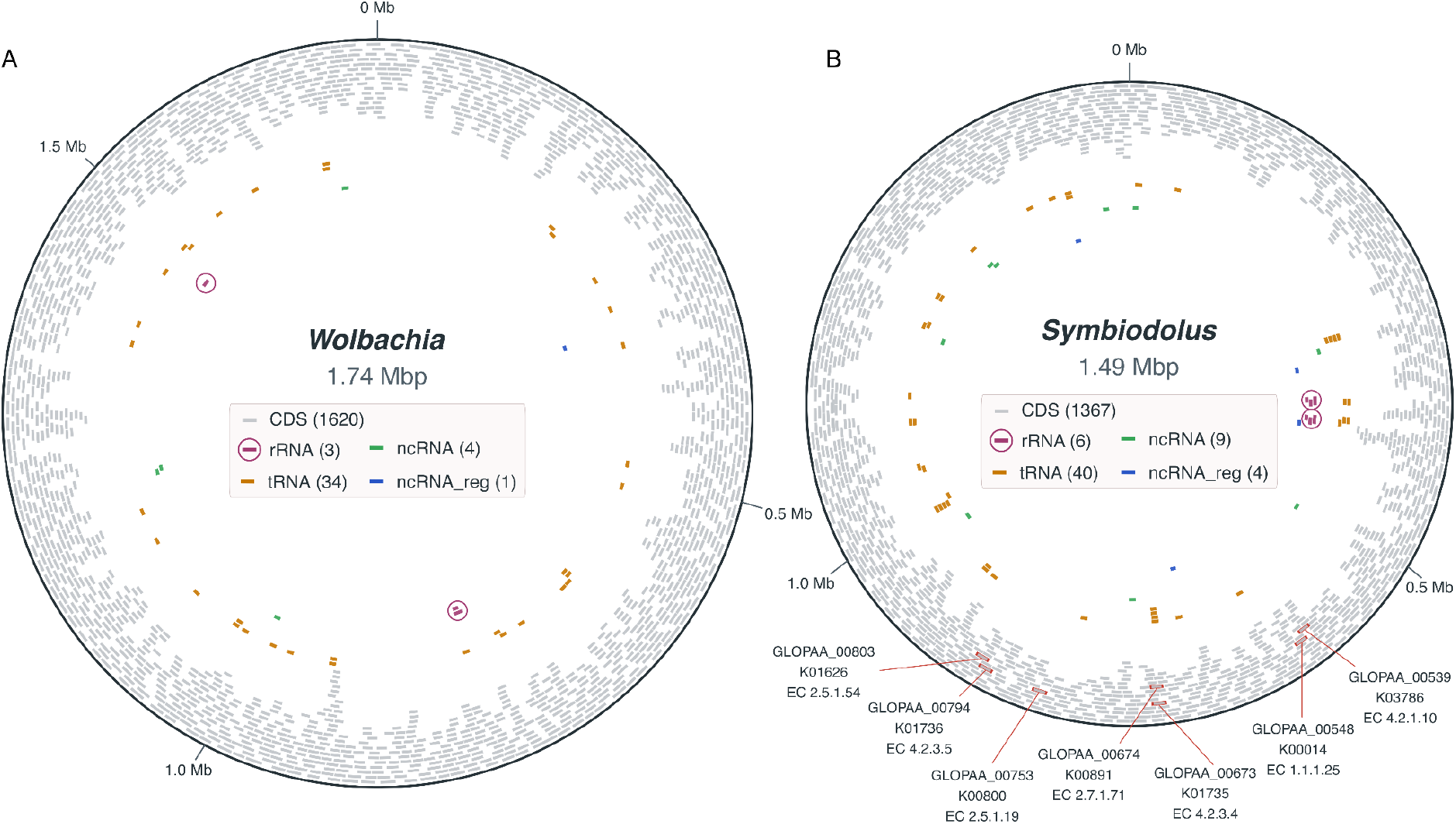
Detected bacterial endosymbionts of *Thymogethes norvegicus*. Schematic circular genome maps showing the distribution of annotated genomic features in **A)** *Wolbachia* and **B)** *Symbiodolus*. Genome sizes are shown at the centers, with scale markings at 0.5-Mbp intervals. Bakta-annotated coding sequences (CDS) are shown closest to the outer circle (in light grey), while transfer RNA (tRNA), ribosomal RNA (rRNA), non-coding RNA (ncRNA), and ncRNA regulatory regions (ncRNA_reg) are shown in distinct bands with distinct colors. Feature counts are provided in the legends. Circles indicate loci containing one to three rRNA genes (16S, 23S, and/or 5S). Shikimate pathway genes were identified from BlastKOALA KEGG Orthology (KO) assignments linked to Bakta locus tags and are labeled (red outline) with their locus tags, KO identifiers, and Enzyme Commission (EC) numbers. Feature centers correspond to annotated genomic midpoints, but segment lengths are rescaled from 71–10,329 bp to 7,500–22,500 bp for visibility and therefore do not represent actual feature lengths.

KEGG pathway analysis identified a complete predicted shikimate pathway leading to chorismate in *Symbiodolus*, but not in *Wolbachia* (Figure 3B; Supplementary Figure 3). Neither endosymbiont had a complete predicted pathway for tyrosine biosynthesis.

## Discussion

The chromosome-level genome assemblies of *Thymogethes norvegicus* show high overall completeness and sequence accuracy. A total of 96.3% and 96.8% of the assemblies were assigned to chromosomes in hap1 and hap2, respectively. BUSCO completeness was 96.6% for hap1 and 90.7% for hap2, while the combined k-mer completeness was 98.1%. Together with the high consensus quality and Hi-C contact maps, these results support the reliability of the assemblies as genomic references.

Although both individuals used for genome assembly, icThyNorv2 (HiFi) and icThyNorv1 (Hi-C), were male, no Y chromosome was identified in either haplotype assembly. Beetles exhibit considerable variation in sex chromosome systems, including both XY and X0 systems, with substantial divergence and degeneration of Y chromosomes (Bracewell et al., 2024). The X chromosome showed sequencing coverage comparable to that of the autosomes, consistent with a single copy of the X chromosome in the male. Whether *T. norvegicus* has an X0 sex-determination system or possesses a Y chromosome that was not identified remains unresolved.

Two bacterial endosymbiont genomes recovered from *T. norvegicus* were identified by 16S rRNA similarity as *Wolbachia* and *Symbiodolus clandestinus*. The *Wolbachia* endosymbiont is likely supergroup A based on ftsZ sequence similarity (Wang et al., 2020; Werren et al., 1995). Bakta annotation and KEGG pathway reconstruction revealed a complete predicted shikimate pathway to chorismate in *Symbiodolus*, but not in *Wolbachia*. Neither possessed a complete predicted pathway for tyrosine biosynthesis. These findings agree with previous reports of chorismate biosynthesis in *Symbiodolus* (Wierz et al., 2024) and shikimate pathway loss in beetle-associated *Wolbachia* (Kiefer et al., 2022). Chorismate production may provide precursors for host cuticle formation, although this nutritional role remains unconfirmed (Wierz et al., 2024).

The chromosome-level genome assembly of *T. norvegicus* provides a foundation for integrating genomic information into future conservation efforts. Population genomic studies could reveal patterns of genetic diversity, population structure, connectivity, and inbreeding among the remaining populations. Such information would be particularly valuable for guiding the selection of source populations for captive breeding, reintroduction, and translocation, which may become increasingly important as local populations continue to decline (Elven, 2024a, 2024b; Røsok, 2024). The reference genome also enables future monitoring of genetic diversity and the effects of conservation interventions, complementing ongoing efforts to protect and restore the habitats of *T. norvegicus* and its host plant, *Dracocephalum ruyschiana* - for which we have also generated a reference genome (Strand et al., 2026).

## Acknowledgements

This project received data management and infrastructure support from ELIXIR Norway, supported by the Research Council of Norway’s grants 322392 and 350529, the University of Bergen, the University of Oslo, the Arctic University of Norway in Tromsø, the Norwegian University of Science and Technology and the Norwegian University of Life Sciences: NMBU. The authors acknowledge support from the National Infrastructure for High Performance Computing and resources provided by Sigma2 as well as Data Storage in Norway (project NN8013K) for computational work. The Norwegian Sequencing Centre generated the sequencing data used in this project (http://sequencing.uio.no).

## Data Availability

Data generated for this study are available under ENA BioProject PRJEB129340. PacBio HiFi sequencing data from specimen icThyNorv2 (ENA BioSample: SAMEA124439599) are deposited under ERR18092818, while Illumina Hi-C sequencing data from specimen icThyNorv1 (ENA BioSample: SAMEA124437175) are deposited under ERR18092792 and ERR18092796. The haplotype-resolved genome assemblies are available under accessions GCA_990133344 (hap1) and GCA_990133343 (hap2).

The genome assemblies and gene annotations for both the host and endosymbionts are available at Zenodo: https://doi.org/10.5281/zenodo.23765747

## References

Artsdatabanken. (2021). Norsk rødliste for arter 2021. https://lister.artsdatabanken.no/rodlisteforarter/2021

Astashyn, A., Tvedte, E. S., Sweeney, D., Sapojnikov, V., Bouk, N., Joukov, V., Mozes, E., Strope, P. K., Sylla, P. M., Wagner, L., Bidwell, S. L., Brown, L. C., Clark, K., Davis, E. W., Smith-White, B., Hlavina, W., Pruitt, K. D., Schneider, V. A., & Murphy, T. D. (2024). Rapid and sensitive detection of genome contamination at scale with FCS-GX. Genome Biology, 25(1), 60.

Bracewell, R., Tran, A., Chatla, K., & Bachtrog, D. (2024). Sex and neo-sex chromosome evolution in beetles. PLoS Genetics, 20(11), e1011477.

Brůna, T., Li, H., Guhlin, J., Honsel, D., Herbold, S., Stanke, M., Nenasheva, N., Ebel, M., Gabriel, L., & Hoff, K. J. (2023). Galba: genome annotation with miniprot and AUGUSTUS. BMC Bioinformatics, 24(1), 327.

Buchfink, B., Xie, C., & Huson, D. H. (2015). Fast and sensitive protein alignment using DIAMOND. Nature Methods, 12(1), 59–60.

Cheng, H., Concepcion, G. T., Feng, X., Zhang, H., & Li, H. (2021). Haplotype-resolved de novo assembly using phased assembly graphs with hifiasm. Nature Methods, 18(2), 170–175.

Chklovski, A., Parks, D. H., Woodcroft, B. J., & Tyson, G. W. (2023). CheckM2: a rapid, scalable and accurate tool for assessing microbial genome quality using machine learning. Nature Methods, 20(8), 1203–1212.

Direktoratet for naturforvaltning. (2010). Handlingsplan for dragehode Dracocephalum ruyschiana og dragehodeglansbille Meligethes norvegicus (p. 56). Direktoratet for naturforvaltning.

Elven, H. (2024a). Inventering og utsetting av dragehodeglansbille i 2023 (p. 33). Naturhistorisk museum, Universitetet i Oslo. https://www.nhm.uio.no/forskning/ressurser/publikasjoner/nhm-rapporter/nhm-rapport-125-2024.pdf

Elven, H. (2024b). Reinventering av dragehodeglansbille i 2022 (p. 83). Naturhistorisk museum, Universitetet i Oslo. https://www.nhm.uio.no/forskning/ressurser/publikasjoner/nhm-rapporter/nhm-rapport-124-2024.pdf

Elven, H., & Pavels, H. (2021). Søk etter dragehodeglansbille Meligethes norvegicus på Hadeland i 2020 (p. 23). Statsforvalteren i Oslo og Viken. https://kudos.dfo.no/documents/47971/files/30396.pdf

Formenti, G., Abueg, L., Brajuka, A., Brajuka, N., Gallardo-Alba, C., Giani, A., Fedrigo, O., & Jarvis, E. D. (2022). Gfastats: conversion, evaluation and manipulation of genome sequences using assembly graphs. Bioinformatics (Oxford, England), 38(17), 4214–4216.

Girgis, H. Z. (2015). Red: an intelligent, rapid, accurate tool for detecting repeats de-novo on the genomic scale. BMC Bioinformatics, 16(1), 227.

Haas, B. J., Salzberg, S. L., Zhu, W., Pertea, M., Allen, J. E., Orvis, J., White, O., Buell, C. R., & Wortman, J. R. (2008). Automated eukaryotic gene structure annotation using EVidenceModeler and the Program to Assemble Spliced Alignments. Genome Biology, 9(1), R7.

Hoff, K. J., & Stanke, M. (2019). Predicting genes in single genomes with AUGUSTUS. Current Protocols in Bioinformatics, 65(1), e57.

Holst, F., Bolger, A. M., Kindel, F., Günther, C., Maß, J., Triesch, S., Kiel, N., Saadat, N., Ebenhöh, O., Usadel, B., Schwacke, R., Weber, A. P. M., Bolger, M. E., & Denton, A. K. (2025). Helixer: ab initio prediction of primary eukaryotic gene models combining deep learning and a hidden Markov model. Nature Methods. 10.1038/s41592-025-02939-1

Jones, P., Binns, D., Chang, H.-Y., Fraser, M., Li, W., McAnulla, C., McWilliam, H., Maslen, J., Mitchell, A., Nuka, G., Pesseat, S., Quinn, A. F., Sangrador-Vegas, A., Scheremetjew, M., Yong, S.-Y., Lopez, R., & Hunter, S. (2014). InterProScan 5: genome-scale protein function classification. Bioinformatics (Oxford, England), 30(9), 1236–1240.

Kanehisa, M., Sato, Y., & Kawashima, M. (2022). KEGG mapping tools for uncovering hidden features in biological data. Protein Science, 31(1), 47–53.

Kanehisa, M., Sato, Y., & Morishima, K. (2016). BlastKOALA and GhostKOALA: KEGG tools for functional characterization of genome and metagenome sequences. Journal of Molecular Biology, 428(4), 726–731.

Kiefer, J. S. T., Schmidt, G., Krüsemer, R., Kaltenpoth, M., & Engl, T. (2022). Wolbachia causes cytoplasmic incompatibility but not male-killing in a grain pest beetle. Molecular Ecology, 31(24), 6570–6587.

Kokot, M., Dlugosz, M., & Deorowicz, S. (2017). KMC 3: counting and manipulating k-mer statistics. Bioinformatics (Oxford, England), 33(17), 2759–2761.

Laetsch, D. R., & Blaxter, M. L. (2017). BlobTools: Interrogation of genome assemblies. F1000Research, 6, 1287.

Li, H. (2013). Aligning sequence reads, clone sequences and assembly contigs with BWA-MEM. In arXiv [q-bio.GN]. arXiv. http://arxiv.org/abs/1303.3997

Li, H. (2018). Minimap2: pairwise alignment for nucleotide sequences. Bioinformatics, 34(18), 3094–3100.

Li, H. (2023). Protein-to-genome alignment with miniprot. Bioinformatics (Oxford, England), 39(1), btad014.

Li, H., Handsaker, B., Wysoker, A., Fennell, T., Ruan, J., Homer, N., Marth, G., Abecasis, G., Durbin, R., & 1000 Genome Project Data Processing Subgroup. (2009). The Sequence Alignment/Map format and SAMtools. Bioinformatics (Oxford, England), 25(16), 2078–2079.

Manni, M., Berkeley, M. R., Seppey, M., Simão, F. A., & Zdobnov, E. M. (2021). BUSCO update: Novel and streamlined workflows along with broader and deeper phylogenetic coverage for scoring of eukaryotic, prokaryotic, and viral genomes. Molecular Biology and Evolution, 38(10), 4647–4654.

Norling, M., Jareborg, N., & Dainat, J. (2018). EMBLmyGFF3: a converter facilitating genome annotation submission to European Nucleotide Archive. BMC Research Notes, 11(1), 584.

Oatley, G., Denton, A., Kovalevskaia, A., & Howard, C. (2025). Sanger Tree of Life HMW DNA Extraction: Manual MagAttract v3. protocols.io. 10.17504/protocols.io.n2bvjedr5gk5/v1

Olberg, S., & Lønnve, O. J. (2021). Kartlegging av dragehodeglansbille i 2021 (p. 24). Stiftelsen Biofokus. https://lager.biofokus.no/biofokus-rapport/biofokusrapport2021-022.pdf

Olberg, S., & Lønnve, O. J. (2022). Bestandsestimering og kartlegging av dragehodeglansbille, Meligethes norvegicus (p. 20). Stiftelsen Biofokus. https://lager.biofokus.no/biofokus-rapport/biofokusrapport2022-130.pdf

Olberg, S., & Lønnve, O. J. (2024a). Kartlegging av dragehodeglansbille i 2024 (p. 16). Stiftelsen Biofokus. https://lager.biofokus.no/biofokus-rapport/biofokusrapport2024-140.pdf

Olberg, S., & Lønnve, O. J. (2024b). Kartlegging og bestandsestimering av dragehodeglansbille i 2023 (p. 26). Stiftelsen Biofokus. https://lager.biofokus.no/biofokus-rapport/biofokusrapport2024-017.pdf

Ranallo-Benavidez, T. R., Jaron, K. S., & Schatz, M. C. (2020). GenomeScope 2.0 and Smudgeplot for reference-free profiling of polyploid genomes. Nature Communications, 11(1), 1432.

Rhie, A., Walenz, B. P., Koren, S., & Phillippy, A. M. (2020). Merqury: reference-free quality, completeness, and phasing assessment for genome assemblies. Genome Biology, 21(1), 245.

Røsok, ø. (2024). Statusrapport for handlingsplan for dragehode og dragehodeglansbille (p. 122).

Statsforvalteren iøstfold, Buskerud, Oslo og Akershus. https://www.statsforvalteren.no/siteassets/fm-oslo-og-viken/miljo-og-klima/rapporter/miljovernavdelingen-i-oslo-og-viken-rapportserie/2024-04-statusrapport-for-handlingsplan-for-dragehode-og-dragehodeglansbille.pdf

Schwengers, O., Jelonek, L., Dieckmann, M. A., Beyvers, S., Blom, J., & Goesmann, A. (2021). Bakta: rapid and standardized annotation of bacterial genomes via alignment-free sequence identification. Microbial Genomics, 7(11). 10.1099/mgen.0.000685

Shaw, J., Marin, M. G., & Li, H. (2026). High-resolution metagenome assembly for modern long reads with myloasm. Nature Biotechnology. 10.1038/s41587-026-03053-z

Shen, W., Sipos, B., & Zhao, L. (2024). SeqKit2: A Swiss army knife for sequence and alignment processing. iMeta, 3(3), e191.

Sim, S. B., Corpuz, R. L., Simmonds, T. J., & Geib, S. M. (2022). HiFiAdapterFilt, a memory efficient read processing pipeline, prevents occurrence of adapter sequence in PacBio HiFi reads and their negative impacts on genome assembly. BMC Genomics, 23(1), 157.

Stanke, M., Schöffmann, O., Morgenstern, B., & Waack, S. (2006). Gene prediction in eukaryotes with a generalized hidden Markov model that uses hints from external sources. BMC Bioinformatics, 7(1), 62.

Strand, M. A., Evju, M., Kyrkjeeide, M. O., Tørresen, O. K., Danneels, B., Arnyasi, M., Ferrari, G., Skage, M., Tooming-Klunderud, A., & Jakobsen, K. S. (2026). A chromosome-level genome assembly of northern dragonhead, Dracocephalum ruyschiana L., 1753.

Tegenfeldt, F., Kuznetsov, D., Manni, M., Berkeley, M., Zdobnov, E. M., & Kriventseva, E. V. (2025). OrthoDB and BUSCO update: annotation of orthologs with wider sampling of genomes. Nucleic Acids Research, 53(D1), D516–D522.

UniProt Consortium. (2023). UniProt: The universal protein knowledgebase in 2023. Nucleic Acids Research, 51(D1), D523–D531.

Wang, X., Xiong, X., Cao, W., Zhang, C., Werren, J. H., & Wang, X. (2020). Phylogenomic analysis of Wolbachia strains reveals patterns of genome evolution and recombination. Genome Biology and Evolution, 12(12), 2508–2520.

Werren, J. H., Zhang, W., & Guo, L. R. (1995). Evolution and phylogeny of Wolbachia: reproductive parasites of arthropods. Proceedings. Biological Sciences, 261(1360), 55–63.

Wierz, J. C., Dirksen, P., Kirsch, R., Krüsemer, R., Weiss, B., Pauchet, Y., Engl, T., & Kaltenpoth, M. (2024). Intracellular symbiont Symbiodolus is vertically transmitted and widespread across insect orders. The ISME Journal, 18(1), wrae099.

Zhang, Z., Schwartz, S., Wagner, L., & Miller, W. (2000). A greedy algorithm for aligning DNA sequences. Journal of Computational Biology: A Journal of Computational Molecular Cell Biology, 7(1–2), 203–214.

Zhou, C., Brown, M., Blaxter, M., Darwin Tree of Life Project Consortium, McCarthy, S. A., & Durbin, R. (2025). Oatk: a de novo assembly tool for complex plant organelle genomes. Genome Biology, 26(1), 235.

Zhou, C., McCarthy, S. A., & Durbin, R. (2023). YaHS: yet another Hi-C scaffolding tool. Bioinformatics (Oxford, England), 39(1), btac808.

