## Supplementary for "A chromosome-level genome assembly of the endangered northern dragonhead sap beetle, *Thymogethes norvegicus* (Easton, 1959)"

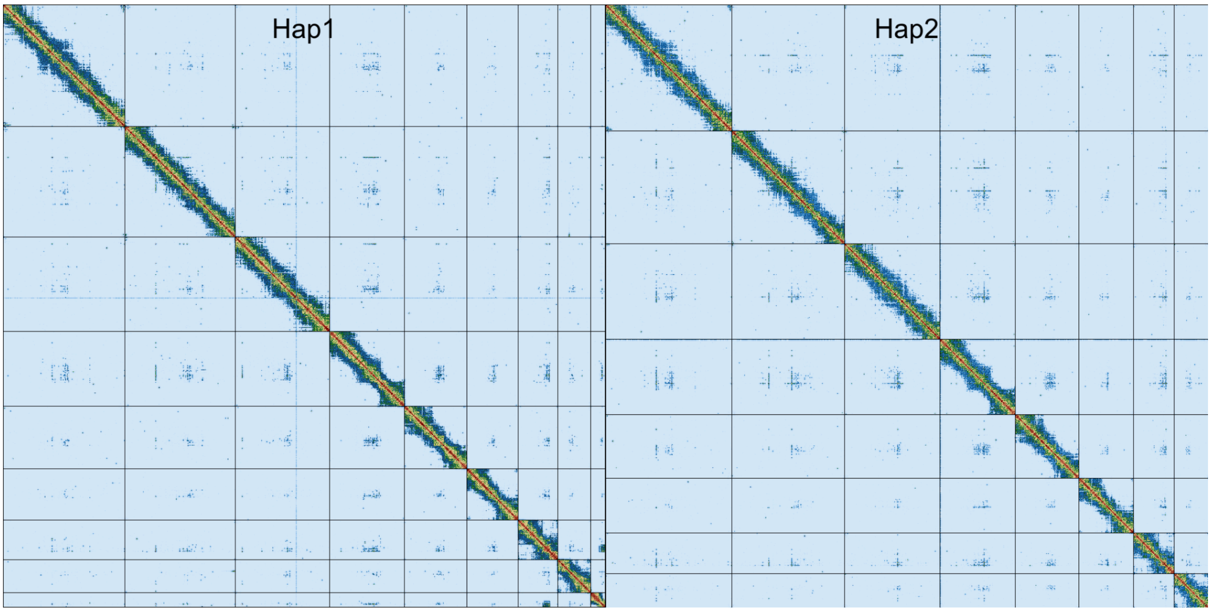


**Supplementary Figure 1. Hi-C contact maps for Thymogethes norvegicus hap1 and hap2.** The contact map displays interaction frequencies between genomic regions, where darker shades represent a higher number of Hi-C contacts. The axes correspond to the coordinates along each assembly. The Hi-C contact maps were generated by mapping the Hi-C reads to the haplotype genomes using BWA-mem, generating a contact map using PretextMap and visualized using PretextSnapshot.

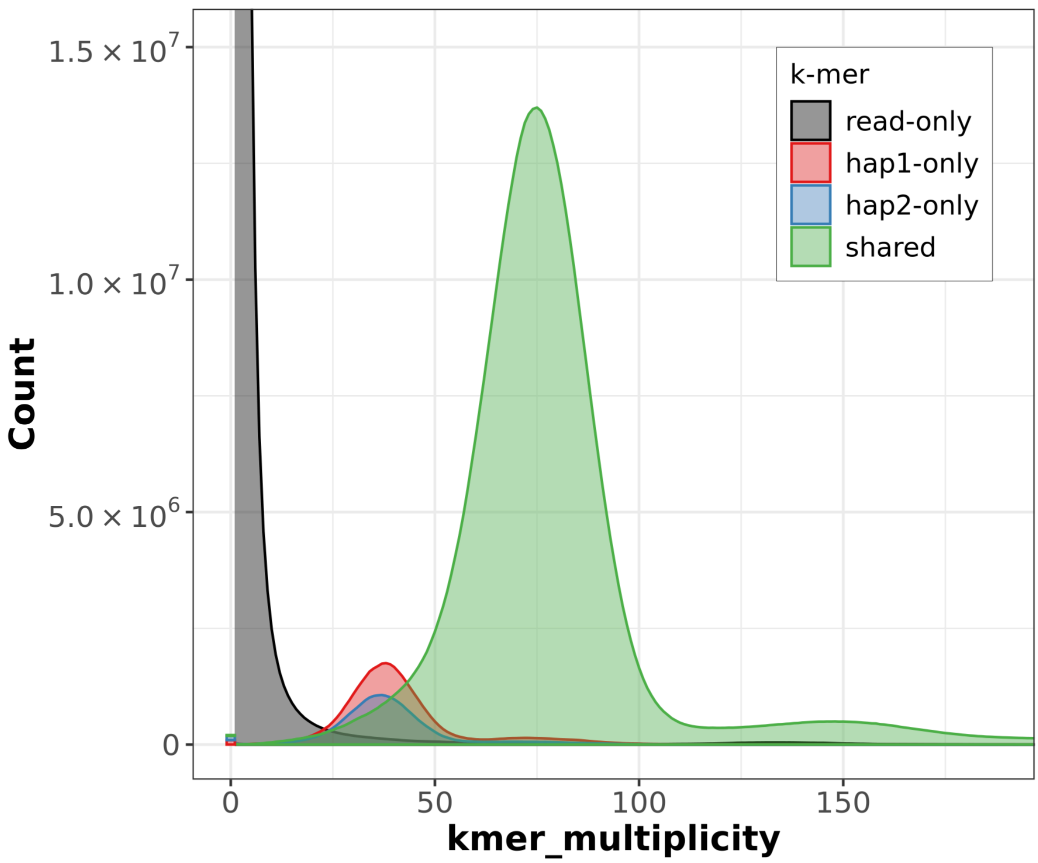


**Supplementary Figure 2. *K*-mer spectra of HiFi reads for Thymogethes norvegicus.** Distributions of *k*-mers found only in the reads (black), only in haplotype 1 (red), only in haplotype 2 (blue), or in both haplotypes (green). The y-axis represents the number of unique *k*-mers, while the x-axis represents the *k*-mer multiplicity (how often the *k*-mer is found in the set of HiFi reads).

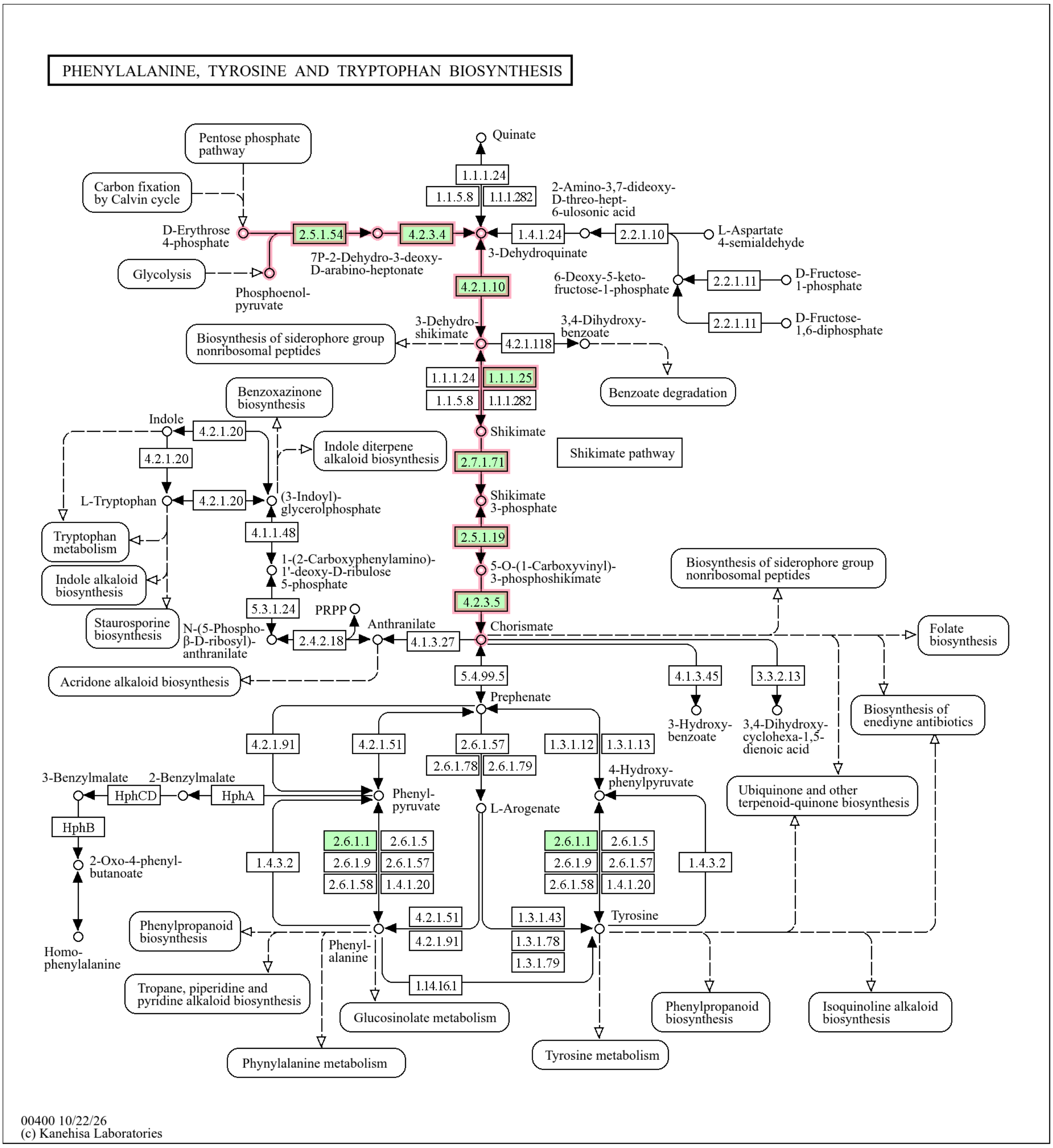


**Supplementary Figure 3. Shikimate pathway in Symbiodolus.** KEGG pathway map of phenylalanine, tyrosine, and tryptophan biosynthesis (map00400), showing enzyme annotations identified through BlastKOALA KEGG Orthology (KO) assignments. The shikimate pathway (module M00022), leading to chorismate, is outlined in red, while green boxes indicate detected enzymes with corresponding KO assignments in Symbiodolus. Enzyme Commission (EC) numbers are shown inside the boxes. The corresponding genes and their genomic positions are shown in Figure 3B. From KEGG Mapper (© Kanehisa Laboratories). Reproduced with permission. Further reproduction requires permission from Kanehisa Laboratories.
